# Neutrophil extracellular traps offer a new therapeutic target for elephant endotheliotropic herpes hemorrhagic disease (EEHV-HD)

**DOI:** 10.1101/2025.08.12.669781

**Authors:** Lisa M. Abegglen, Aaron Rogers, Gareth Mitchell, C. Bradley Nelson, Madison I. Sanborn, Ryan Kennington, McKenna Rogers, Virginia R. Pearson, Miranda Sharp, Lauren L. Howard, Erin Latimer, Jennifer A. Landolfi, Christine Molter, Erika Crook, Wendy Kiso, Dennis Schmitt, Paul D. Ling, Kimberly Martinod, Joshua D. Schiffman

**Affiliations:** Department of Pediatrics, University of Utah, Salt Lake City, Utah, USA; Hunstman Cancer Institute, University of Utah, Salt Lake City, Utah, USA; Peel Therapeutics, Inc., Salt Lake City, Utah, USA; Program in Blood Cell Development and Function, Fox Chase Cancer Center, Philadelphia, Pennsylvania, USA; North American EEHV Advisory Group; Elephant Herpesvirus Laboratory, Smithsonian’s National Zoo and Conservation Biology Institute, Washington, DC, USA; Zoological Pathology Program, University of Illinois, Brookfield, IL, USA; Houston Zoo, Houston, Texas, USA; Utah’s Hogle Zoo, Salt Lake City, Utah, USA; Center for Applied Reproductive & Emerging Sciences, Fort Worth Zoo, Fort Worth TX, USA; Department of Animal Science, William H. Darr College of Agriculture, Missouri State University, Springfield, MO, USA; Department of Molecular Virology and Microbiology, Baylor College of Medicine, Houston, Texas, USA; Aab Cardiovascular Research Institute, Department of Medicine, University of Rochester School of Medicine and Dentistry, Rochester, New York, USA

**Author notes:** Correspondence should be directed to Dr. Abegglen.

## Abstract

Elephant survival is threatened by a devastating hemorrhagic disease called elephant endotheliotropic herpes virus-hemorrhagic disease (EEHV-HD). Once clinical signs are observed in elephants, the disease progresses rapidly and frequently results in death. EEHV-HD negatively impacts elephant conservation because very young, reproductively immature elephants are most at risk for death. Ongoing efforts to understand disease pathogenesis and progression may identify treatment targets and improve clinical outcomes. In some lethal EEHV-HD cases, microthrombosis has been observed in organ tissues similar to other hemorrhagic diseases in humans and animals where sticky webs of protein-coated DNA strands called neutrophil extracellular traps (NETs) exacerbate thrombosis and hemorrhage associated with disseminated intravascular coagulation (DIC). In this study, we sought to identify if NET formation occurs in elephants and could contribute to poor outcomes in EEHV-HD. Our study demonstrated NET release for the first time from elephant heterophils (neutrophils) that occurred in response to various stimuli, including plasma from EEHV-HD affected elephants. EEHV-HD affected tissues contained extensive NETs suggesting that dysregulated NET formation contributes to pathogenesis of this disease. Importantly, elephant neutrophils were blocked from releasing NETs in response to EEHV-HD plasma using known NET inhibitors. The ability to stop NETs in EEHV-HD offers a new therapeutic approach that could be combined with current therapies to improve survival for affected elephants and to positively impact conservation efforts.

## Introduction

Elephant endotheliotropic herpesviruses (EEHV) are viruses endemic to both Asian and African elephants (*Elephas maximus* and *Loxodonta africana*) worldwide^1–9^. Seven species and multiple subtypes of EEHV are known to infect elephants, and all elephants are believed to carry some of these viruses as latent infections ^2,6,10–14^. A properly primed immune system protects elephants from disease when these viruses become activated ^12,15^. However, the immature immune system and loss of maternally derived antibodies in young elephants, as well as EEHV naïve or compromised immune systems of adult elephants can leave them susceptible to rapidly increasing EEHV in circulation (viremia) and to illness or death from disease ^6,16,17^. Exponentially increasing EEHV viremia can cause a highly fatal hemorrhagic disease (EEHV-HD) that primarily affects young, immunologically naive elephants of both species ^3,5,6,9,13,14,18^. EEHV-HD occurs in elephants under human care and free ranging elephants. The endangered status of Asian and African elephants ^19,20^ makes EEHV-HD-related mortality a significant conservation concern.

Clinical signs of EEHV-HD include lethargy, altered mental status, decreased food or water intake, lameness/stiffness, cranial or front limb edema, petechial hemorrhages, and cyanosis of the tongue ^9,13,21^. Severe cases progress rapidly, leading to internal hemorrhaging and organ failure often within 1-7 days after clinical signs of illness are observed ^18^. Not all elephants with EEHV viremia develop EEHV-HD, but for those who do, death is frequently the outcome ^6,7^. Extensive efforts to treat affected elephants have increased survival. Treatments include fluids, antiviral medications, analgesics, and transfusion with plasma from EEHV seropositive elephants ^5,7,14,22,23^. Because each treatment has not been tested in clinical trials, it is challenging to determine which are beneficial. The epidemiology and pathophysiology that influence EEHV viremia and progression to EEHV-HD are currently under global investigation in both free ranging and managed elephant populations ^4,8,24–26^. Despite the need for more knowledge, it is clear from case review that early detection and intervention are crucial for successful treatment outcomes ^7,14,23,27^. Vaccine development is underway, and effective vaccines will protect some elephants from EEHV-HD induced by rapidly increasing viremia and save lives ^28,29^. Even as effective vaccines for both Asian and African elephants become available, a tremendous need exists to better understand disease progression from viremia to fatal hemorrhagic disease. Unvaccinated elephants and elephants with ineffective responses to vaccination will still require effective treatment. With a better understanding of the pathophysiology leading to severe disease and death, opportunities for therapeutic intervention may become clear.

In the context of viral infections, sepsis can occur due to the body’s overwhelming immune response leading to widespread inflammation and organ dysfunction ^30^. Evidence suggests that sepsis occurs in the context of EEHV-HD ^31,32^. Like sepsis observed in other species, elephants with severe cases of EEHV-HD develop hypotension, tachycardia, and altered mental status ^21,33^. Additionally, a recent study found increased expression of pro-inflammatory cytokines in EEHV-HD affected tissues ^32^ and evidence of disseminated intravascular coagulopathy (DIC) ^32^, an end-stage phenomenon also observed in humans with sepsis ^34^. EEHVs infect capillary endothelial cells, and virus-mediated endothelial damage leads to increased vascular permeability and leakage with severe widespread edema and hemorrhage ^32,35^. Vascular compromise is known to influence the development of sepsis in other diseases ^36^. In human patients, the release of pro-inflammatory cytokines and activation of coagulation pathways further exacerbate systemic inflammation and microvascular thrombosis in DIC ^37^. Early interventions used to treat EEHV-HD include treatments with efficacy against sepsis in human patients such as fluid resuscitation, vasopressor support, antibiotics to target secondary bacterial infections, and transfusion of blood products to address coagulopathy ^38^.

Understanding the role of dysregulated immune responses in the pathogenesis of EEHV-HD remains crucial for implementing effective management strategies and improving survival rates in affected elephants. Discovery of underlying mechanisms that contribute to DIC and microthrombi in EEHV-HD may reveal novel targets for therapeutic interventions to mitigate EEHV-HD’s fatal impact on elephants and conservation. In humans, dysregulated neutrophil responses are known to exacerbate virus induced disease and contribute to death ^39,40^. When neutrophils are activated, they can release neutrophil extracellular traps (NETs). NETs are webs of decondensed chromatin with attached enzymes to capture and degrade pathogens ^41^. While NETs play a beneficial role as a first response to control infection, dysregulated NET release contributes to the pathophysiology of many diseases ^41^. For example, neutrophils can be activated to form NETs in response to a variety of stimuli including pathogens, immune activation, endothelial cell damage, activated platelets, and coagulation factors leading to induction of clotting, which can further damage endothelial cells and exacerbate bleeding ^42,43^. We previously described a role for NETs in the pathogenesis of COVID-19 in humans, including microthrombosis associated with COVID-19 induced acute respiratory distress syndrome (ARDS) ^40^. EEHV-HD causes widespread hemorrhage and coagulopathy with increased microthrombosis,^31,32,35^ a histologic finding suggestive of NET-mediated disease. Here, we identified increased NET formation in EEHV-HD affected tissues and demonstrated the ability of elephant heterophils (functionally equivalent to neutrophils ^15^) to release NETs. Our description of NETs for the first time in elephant cells, including their presence in EEHV-HD affected tissues, expands our knowledge of the pathological mechanisms involved in this devastating disease. We introduce elephant NETs as a new therapeutic target to potentially improve outcomes in affected elephants.

## Methods

### Neutrophil isolation

Whole blood was collected from Asian and African elephants housed at Houston Zoo and Utah’s Hogle Zoo (respectively) by venipuncture into EDTA-coated vacutainer tubes under approval from each institution’s internal science review committee. Blood was processed to isolate elephant heterophils within 30 minutes of collection. Tubes were placed in an incubator for 15-30 minutes for red blood cells (RBCs) sedimentation. Overlying suspension containing white blood cells (WBCs) was collected and transferred to a fresh tube. PBS was added and the tube was centrifuged at 1500rpm for five minutes with no break to pellet the cells. The cells were resuspended in media (RPMI with 0.05% FBS heat inactivated at 70°C for 30 minutes) and layered onto Mono-Poly Resolving Medium (MP Biomedicals) followed by centrifuging for 30 minutes at 300 x g in a swinging bucket rotor at room temperature with the brake off to allow proper separation. The layer of cells containing heterophils was harvested using a pipette, avoiding contamination from the other layers.

### In vitro neutrophil/heterophil extracellular trap formation assays

For confocal imaging, 8 well glass chamber slides were coated with poly-L-lysine. Half a million cells were seeded in each well in RPMI media with 0.05% heat inactivated FBS. FBS was inactivated at 70°C for 30 minutes to inactivate DNase in serum. Cells were stimulated with the indicated concentrations of PMA (phorbol 12-myristate 13-acetate), LPS (lipopolysaccharide), ionomycin, or EEHV-HD positive plasma for 2 hours at 37C. The cells were then fixed and stained for imaging. For inhibition experiments, cells were pretreated with the indicated concentrations of inhibitors (dexamethasone from Selleck Chemicals S1322 or BMS-P5 from Cayman Chemical 33581) for 30 minutes prior to adding stimulants at the indicated concentrations.

For real-time microscopy, MPO-FITC and Sytox AADvanced were added to RPMI media with 0.05% heat inactivated FBS + 0.01M HEPES. PMA, LPS, and Ionomycin were added at the indicated concentrations in a 96 well plate media containing dye (MPO-FITC and SYTOX AADvanced). SYTOX is a DNA stain that labels extracellular DNA and DNA from cells with compromised cell membranes. Elephant heterophils were then stained for viability and counted by Trypan Blue exclusion. Cells were seeded at 10,000 cells/well of the 96 well plate. The plates were then imaged in the Incucyte (Sartorius) every 15 minutes, capturing 2 images per well, at 650ms GFP and 500ms RFP. Data was analyzed by measuring area of overlap for MPO-FITC and SYTOX by phase confluence using Incucyte 2023 Rev2 software.

### Tissue Samples

Tissue samples from Asian Elephants with lethal EEHV-HD and from Asian elephants that died of other causes were collected at necropsy and frozen immediately or fixed in formalin (Table 1). Tissues were sectioned to 5 μm thickness and mounted on slides.

**Table 1:** Asian elephant tissues analyzed and quantified for NET formation.

| Species | Common Name | Age | Sex | Cause of Death | EEHV Type | Heart | Liver | Lung | Brain | Spleen |
| --- | --- | --- | --- | --- | --- | --- | --- | --- | --- | --- |
| Elephas maximus | Asian elephant | 2 | M | EEHV-HD | EEHV1A | + | + | + | + |  |
| Elephas maximus | Asian elephant | 3 | M | EEHV-HD | EEHV1A | + | + | + | + | + |
| Elephas maximus | Asian elephant | 4 | M | EEHV-HD | EEHV1A | + |  |  |  | + |
| Elephas maximus | Asian elephant | 5 | M | EEHV-HD | EEHV1A | + | + | + | + |  |
| Elephas maximus | Asian elephant | 6 | F | EEHV-HD | EEHV1A |  |  |  | + |  |
| Elephas maximus | Asian elephant | 6 | F | EEHV-HD | EEHV1A |  |  | + |  |  |
| Elephas maximus | Asian elephant | 9 | F | EEHV-HD | EEHV1A |  | + | + |  | + |
| Elephas maximus | Asian elephant | 11 | F | EEHV-HD | EEHV1A | + |  | + |  |  |
| Elephas maximus | Asian elephant | 13 | M | EEHV-HD | EEHV1A |  | + | + | + | + |
| Elephas maximus | Asian elephant | 0 | F | Other | NT | + |  |  |  |  |
| Elephas maximus | Asian elephant | 44 | F | Other | NT | + |  |  |  |  |
| Elephas maximus | Asian elephant | 44 | M | Other | NT |  |  | + |  |  |
| Elephas maximus | Asian elephant | 46 | F | Other | NT |  |  | + |  |  |
| Elephas maximus | Asian elephant | 47 | F | Other | NT |  | + | + |  |  |
| Elephas maximus | Asian elephant | 53 | F | Other | NT | + | + | + | + | + |
| Elephas maximus | Asian elephant | 60 | F | Other | NT | + | + |  | + | + |
| Elephas maximus | Asian elephant | 73 | F | Other | NT | + | + |  | + |  |
| Elephas maximus | Asian elephant | Unknown | F | Other | NT |  |  | + |  |  |
| Elephas maximus | Asian elephant | Unknown | F | Other | NT |  |  | + |  |  |

### Immunofluorescence

Tissue samples were stained with rabbit anti-human citrullinated histone H3 (1:400, ab5103; Abcam) and goat anti-human/mouse MPO (1:67, AF3667; R&D Systems) with AlexaFluor488-conjugated anti-rabbit and DyLight 649-conjugated anti-goat secondary antibodies). ProLong Glass Antifade Mountant formulated with NucBlu (Hoechst 33342) was used to stain DNA (P36981; Invitrogen). Images were captured by confocal microscopy (Leica SP8) and analyzed using FIJI/Image-J (version 2.1.0/1.53c; NIH).

### Imaging and Quantification

All samples were deidentified prior to imaging to blind the results during image collection and quantification. First, whole slide immunofluorescence images were collected by confocal microscopy (Leica SP8) by one individual. A second individual then reviewed the slides and selected ten 4-by-4 tiled representative sections with the most citrullinated histone H3 and MPO staining. A third individual then collected high resolution images of each 4-by-4 section by confocal microscopy. Each blinded section was analyzed using FIJI/Image-J’s cell counter plug-in (version 2.1.0/1.53c; NIH) to calculate the percent of citrullinated histone H3 positive cells, relative to DAPI (4’,6-diamindino-2-phenylindole).

## Results

### Elephant heterophils form extracellular traps in response to known NET inducing stimuli

Heterophils (functionally equivalent to neutrophils ^15^) from both Asian and African elephants were enriched from fresh whole blood by red blood cell sedimentation and granulocyte isolation by density gradient centrifugation (**Figure 1A)**. Heterophils comprise 40% of the circulating white blood cells of elephants ^44^, and a similar approach was used to successfully isolate elephant heterophils with high viability (99%) and purity (98%) ^45^. In the current study, heterophil viability was 97% and purity was 79% after enrichment (**Figure 1B**). Processing was initiated within 30 minutes of blood draw due to the short half-life of neutrophils/heterophils ^46^. Attempts to measure NET formation with stimulation failed if blood was not processed within 30 minutes of collection, due to high heterophil death and release of NETs in the untreated control samples. Enriched heterophils were treated with stimuli known to induce NET formation in other species: phorbol 12-myristate 13-acetate (PMA), and lipopolysaccharide (LPS), and ionomycin ^47^. When neutrophils form NETs, chromatin decondenses and histone H3 becomes citrullinated ^48^. To determine if stimulated elephant heterophils form and release NETs, immunofluorescence was performed to measure expression citrullinated histone H3 (citH3). Minimal citH3 expression was observed in unstimulated cells, and citH3 expression increased with increasing doses of stimuli in African (**Figures 2A and 2B**) and Asian elephant heterophils (**Figures 3A and 3B)**. As a secondary measure of NET formation and release from cells, intracellular DNA and extracellular DNA were visualized in blue and yellow, respectively, after African elephant heterophils were treated with ionomycin (**Figure 2C**). Large webs of DNA (NET) were released from heterophils. NET formation increased overtime in a dose dependent manner when African elephant heterophils were treated with PMA, LPS, or ionomycin as measured in a third assay by real-time microscopy for overlap in DNA and myeloperoxidase (MPO), an enzyme in heterophil granules that is attached to the DNA when chromatin decondenses to form NETs (**Figure 2D**). Elephant heterophils formed significantly more NETs in response to stimulation compared to unstimulated heterophils (AUC p<0.0001 for highest two concentrations of stimulants compared to untreated). Given the limitations related to collecting and processing fresh samples and the lack of access to real-time live-cell imaging, detecting citH3 by confocal microscopy was the most practical method for visualizing and quantifying NETs, as it allowed for cell fixation after stimulation.

**Figure 1:**
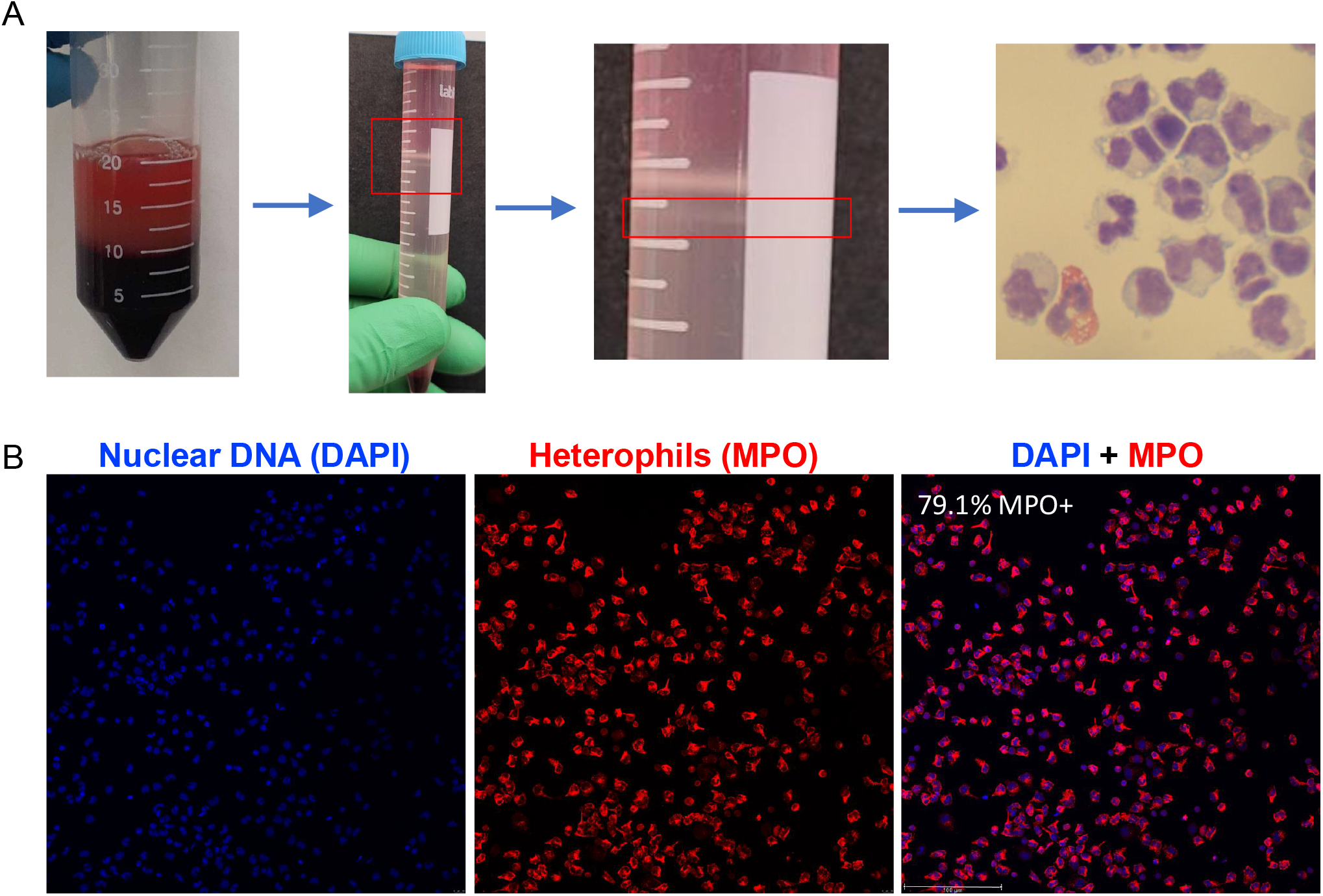
Elephant heterophil isolation. A. Whole venous blood was collected in EDTA tubes. Following red blood cell sedimentation, white blood cells were separated by density gradient centrifugation. The bottom layer (granulocytes) of white blood cells was analyzed by microscopy-images of cytospin preparations. Heterophils were identified by nuclear structure. B. The percent of heterophils in the isolated granulocyte layer was determined by staining for myeloperoxidase (MPO) expression. MPO+ cells/Total cells (DAPI) was 79.1%.

**Figure 2:**
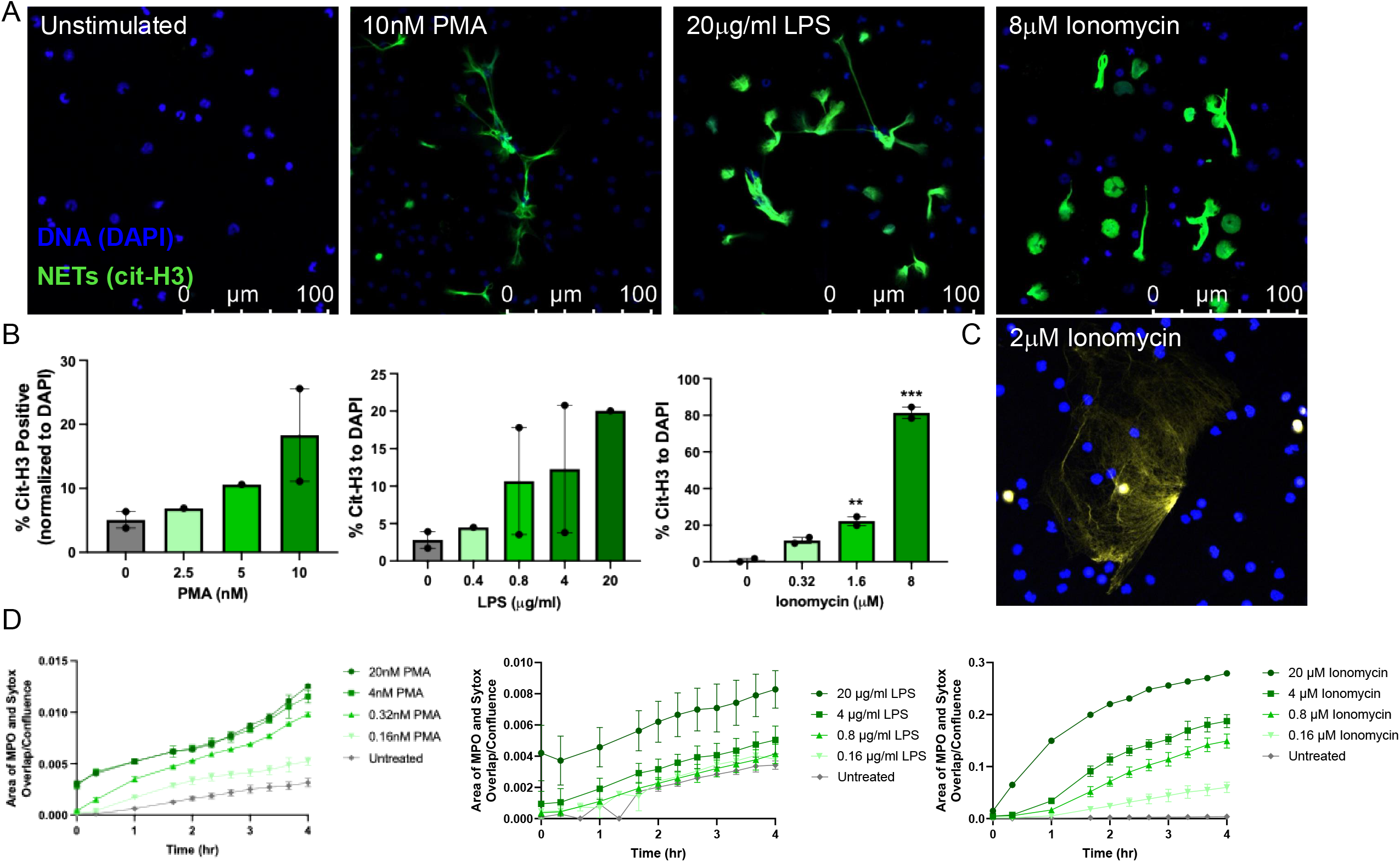
African elephant heterophils formed NETs in response to stimuli. A. Representative images of African elephant heterophils unstimulated, stimulated with 10nM PMA, 20μg/ml LPS or 8μM ionomycin. Nuclear DNA (DAPI) is in blue and citrullinated histone H3 (citH3) is in green. B. Dose dependent response of African elephant heterophils to PMA, LPS, and ionomycin stimulation. Percent of citH3 cells/total cells (DAPI) is graphed. Dots represent replicate experiments performed with blood drawn on different days from two different individuals. C. NET release from an African elephant heterophil stimulated with 2μM ionomycin measured with Sytox Orange (yellow) to label extracellular DNA and DNA in cells with compromised membranes, and Hoescht 33342 to label intracelluar DNA (blue). D. African elephant heterophils responded to increasing doses of stimuli over time as measured by real-time microscopy. Area under the curve (AUC) untreated (UT) vs. 4nM PMA p<0.0001, UT vs. 20mg/ml p<0.0001, UT vs. 4μg/ml LPS p<0.005, UT vs. 20μg/ml LPS p<0.0001, UT vs. 0.16μM ionomycin (IO) p<0.0001, UT vs. 0.8μM IO p<0.0001, UT vs. 4μM IO p<0.0001, UT vs. 20μM IO p<0.0001.

**Figure 3:**
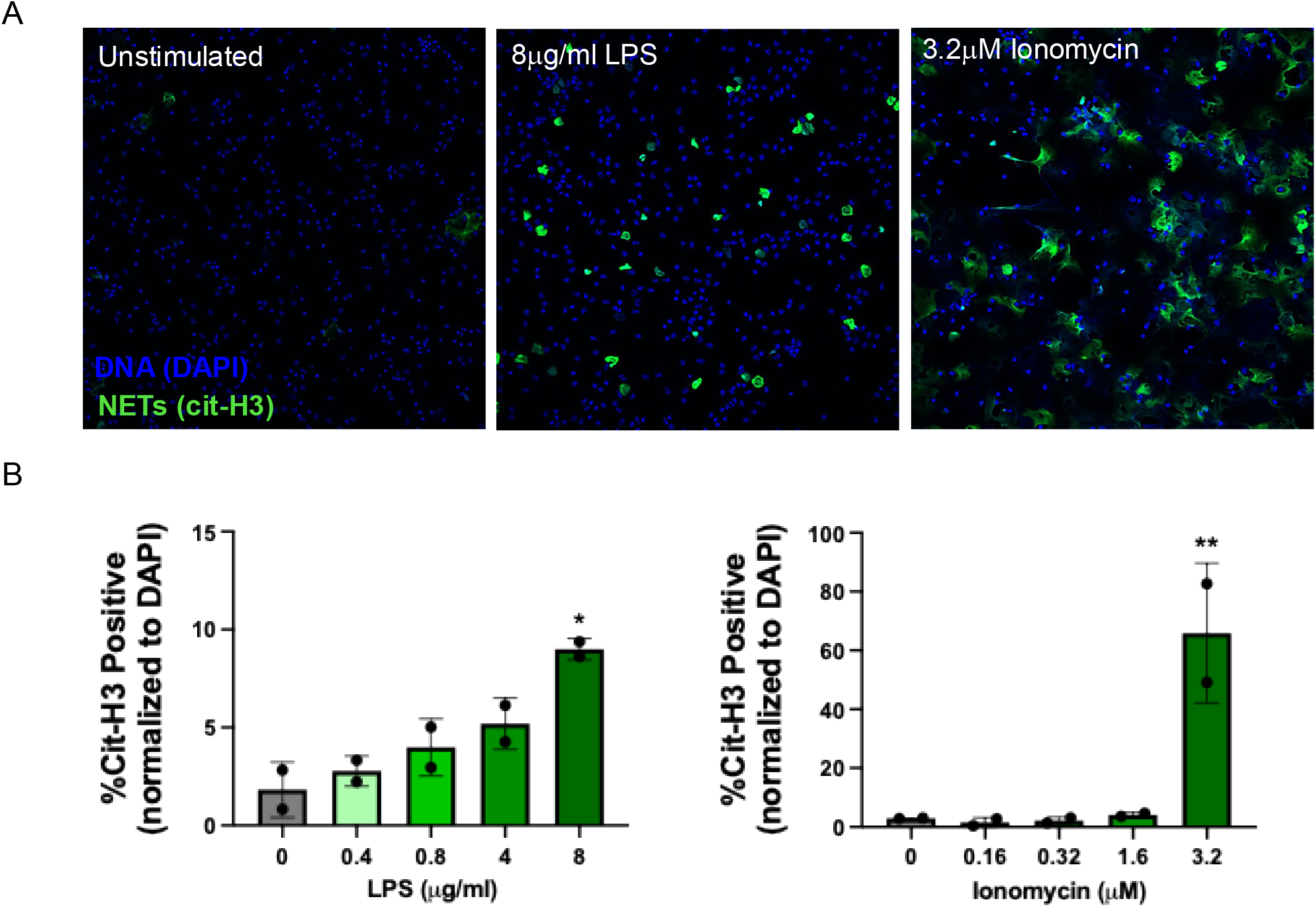
Asian elephant heterophils formed NETs in response to stimuli. A. Representative images of Asian elephant heterophils unstimulated, stimulated with 8μg/ml LPS or 3.2μM ionomycin. Nuclear DNA (DAPI) is in blue and citrullinated histone H3 (citH3) is in green. B. Dose dependent response of Asian elephant heterophils to LPS and ionomycin stimulation. Percent of citH3 cells/total cells (DAPI) is graphed. Dots represent replicate experiments performed with blood drawn on different days from three different individuals.

### Plasma from EEHV-HD affected elephants stimulates NET formation and release

NETs are formed and released from human neutrophils during viral infection ^40,49^. To determine if factors present in the plasma during active EEHV-HD stimulate elephant heterophils to form NETs, plasma was collected from a very sick Asian elephant during active EEHV viremia (confirmed by PCR) with symptoms of EEHV-HD who ultimately died from hemorrhagic disease, determined by the presence of lesions indicative of HD detected histologically in tissues collected at necropsy. When treated with this EEHV-HD positive plasma, healthy Asian elephant heterophils formed and released NETs, while control plasma collected from an Asian elephant negative for EEHV viremia did not stimulate NET formation (**Figure 4A and 4B**). Next, we treated African elephant heterophils with the same EEHV-HD positive plasma and again demonstrated NET formation and release (**Figure 4C and 4D**).

**Figure 4:**
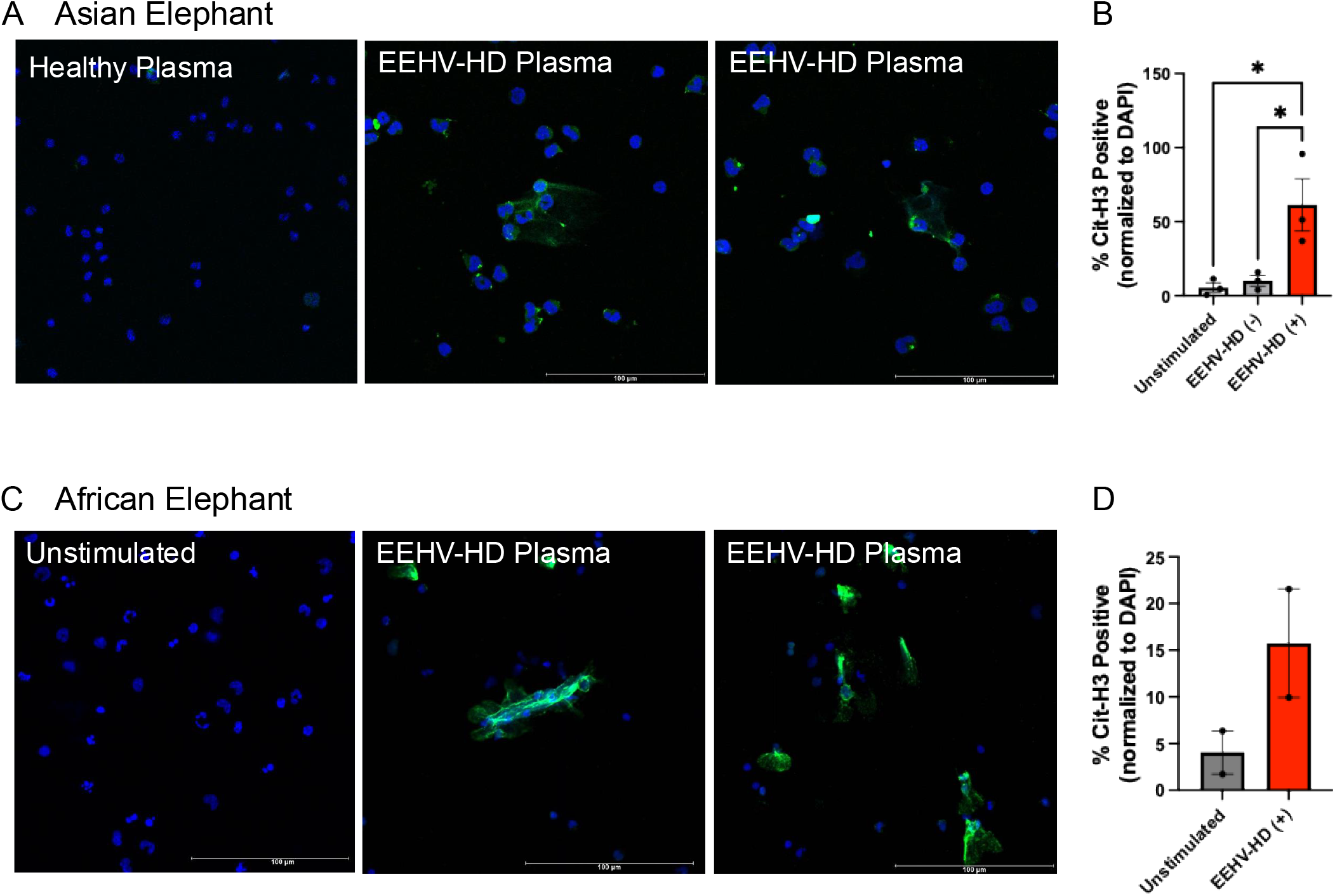
Asian and African elephant heterophils formed NETs in response to plasma collected during active EEHV-HD. A. Representative images and B. quantification of Asian elephant heterophils treated with healthy or EEHV-HD positive plasma (2 example images). C. Representative images and D. quantification of African elephant heterophils untreated or treated with EEHV-HD positive plasma (2 example images). Nuclear DNA (DAPI) is in blue and citrullinated histone H3 (citH3) is in green. Percent of citH3 cells/total cells (DAPI) is graphed. Dots represent replicate experiments performed with blood drawn on different days from different individuals.

### NETs are present in EEHV-HD affected tissues

Disseminated intravascular coagulation (DIC) has been documented as a complication of EEHV-HD ^31^. DIC is characterized by widespread activation of the clotting cascade, resulting in the formation of blood clots throughout the body’s small blood vessels ^50^. This can lead to multiple organ damage and failure due to restricted blood flow, as well as a depletion of clotting factors and platelets that increases the risk of severe bleeding ^51^. Dysregulated NET formation and release plays a significant role in the development and progression of DIC. Excessive formation of NETs contributes to hypercoagulation by providing a scaffold for platelet, von Willebrand factor, and fibrin, which promotes thrombus formation in the microvasculature. NETs are also known to directly damage vascular endothelial cells and amplify inflammation, further exacerbating the coagulopathy seen in DIC ^49,52,53^. To determine if NETs were present in tissues from elephants who succumbed to EEHV-HD, immunofluorescence to detect citrullinated histone H3 was performed on tissues collected at necropsy from elephants whose cause of death was EEHV-HD and elephants who died of other causes. **Table 1** lists the species, sex, age, virus type, and tissue type of the samples tested. Tissue samples from hearts, livers, lungs, spleens, and brains were imaged for the presence of heterophils (MPO) and NETs (citH3) (**Figure 5A**). Quantification of NET-forming cells and released NETs in these tissues revealed a significant increase in NETs in EEHV-HD affected tissues compared to non EEHV-HD tissues (heart p<0.005, liver p<0.05, lung p<0.05, brain p<0.0001, spleen p<0.005) (**Figure 5B**).

**Figure 5:**
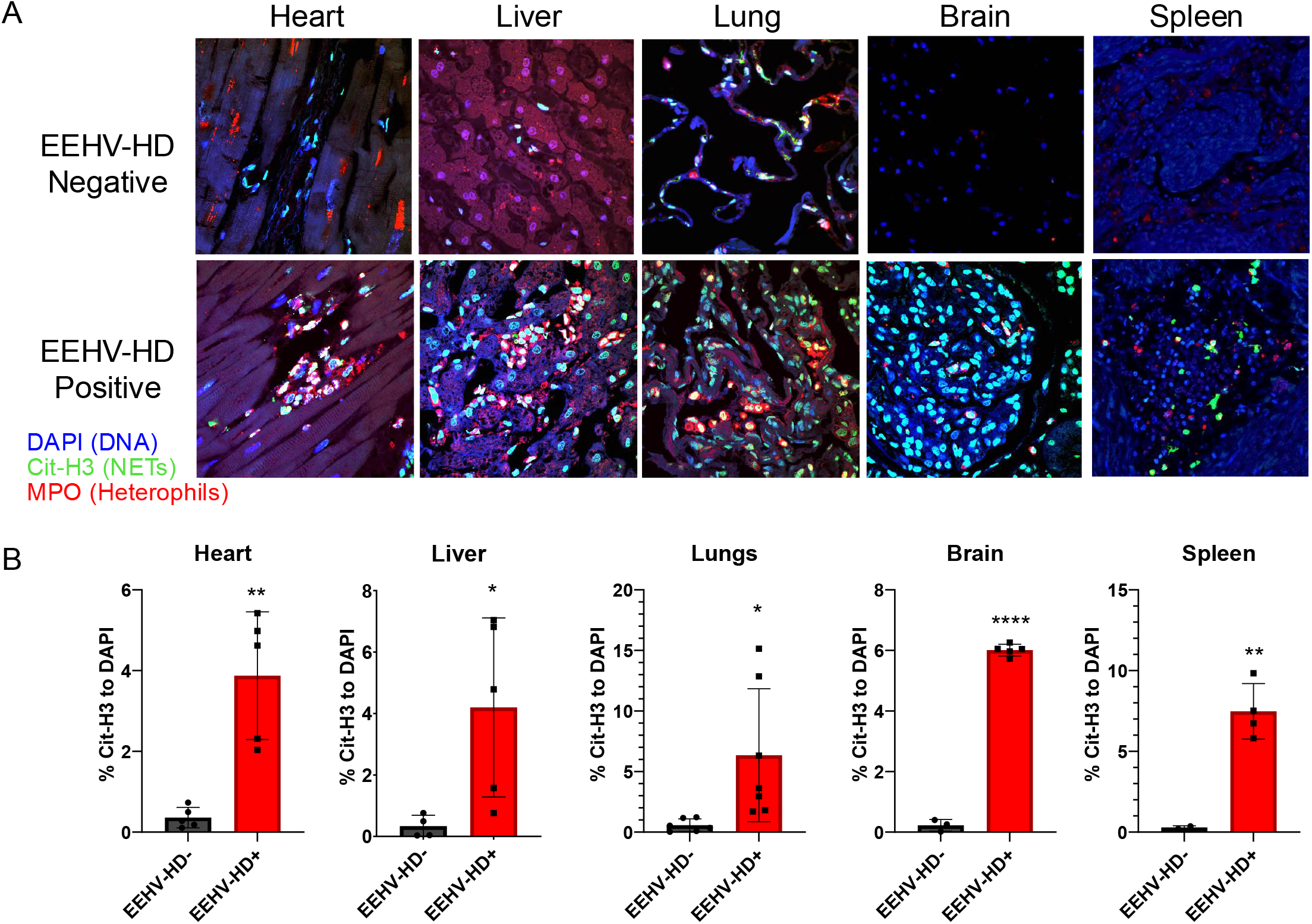
NETs formed in tissues from EEHV-HD affected Asian elephants. A. Representative images of necropsy tissues from Asian elephants that died of EEHV-HD and control elephants that died of other causes. Nuclear DNA is shown in blue (DAPI), heterophils are shown in red (MPO), and NETs are shown in green (citH3). B. Quantification of NET formation in tissues. Each dot represents an individual Asian elephant. * p<0.05, **p<0.005, **** p<0.0001

### NET inhibitors block elephant NET formation in response to stimuli, including EEHV-HD positive plasma

In multiple lethal mouse models of sepsis and viral disease, blocking NET formation or degradation of NETs improved survival ^54–57^. Multiple NET-specific inhibitors are in preclinical and clinical development for human diseases ^58^. BMS-P5 is a compound under clinical development that effectively inhibits NET formation by selectively targeting PAD4, which is an enzyme critical for NET formation ^59^. Another strategy to broadly suppress NET formation is through general immune suppression with steroid drugs like dexamethasone. Dexamethasone inhibits NET formation ^60^ and has been shown to improve outcomes in some human viral diseases, including COVID-19 ^61^. To determine if BMS-P5 or dexamethasone could inhibit elephant NET formation, African and Asian elephant heterophils were treated with these inhibitors prior to stimulating with PMA, ionomycin (IO), or LPS. NET formation was measured by staining for citH3 and DNA. Decreased NET formation was observed in stimulated elephant heterophils that were pretreated with either BMS-P5 or dexamethasone (**Figure 6A and 6B**) compared to those treated with vehicle (PMA compared to BMS-P5 + PMA p<0.0005, PMA compared to dexamethasone + PMA p<0.0005, IO compared to BMS-P5 + IO p<0.01, IO compared to dexamethasone + IO p<0.01, LPS compared to dexamethasone + LPS p<0.05). Next, we measured the ability of these drugs to inhibit NET formation in heterophils stimulated with EEHV-HD positive plasma. BMS-P5 and dexamethasone inhibited NET formation when African or Asian elephant heterophils were stimulated with EEHV-HD positive plasma (**Figure 7A-C**), suggesting that dysregulated NET formation can be inhibited in the context of EEHV-HD (EEHV-HD positive plasma stimulated compared to pretreatment with BMS-P5 p<0.01).

**Figure 6:**
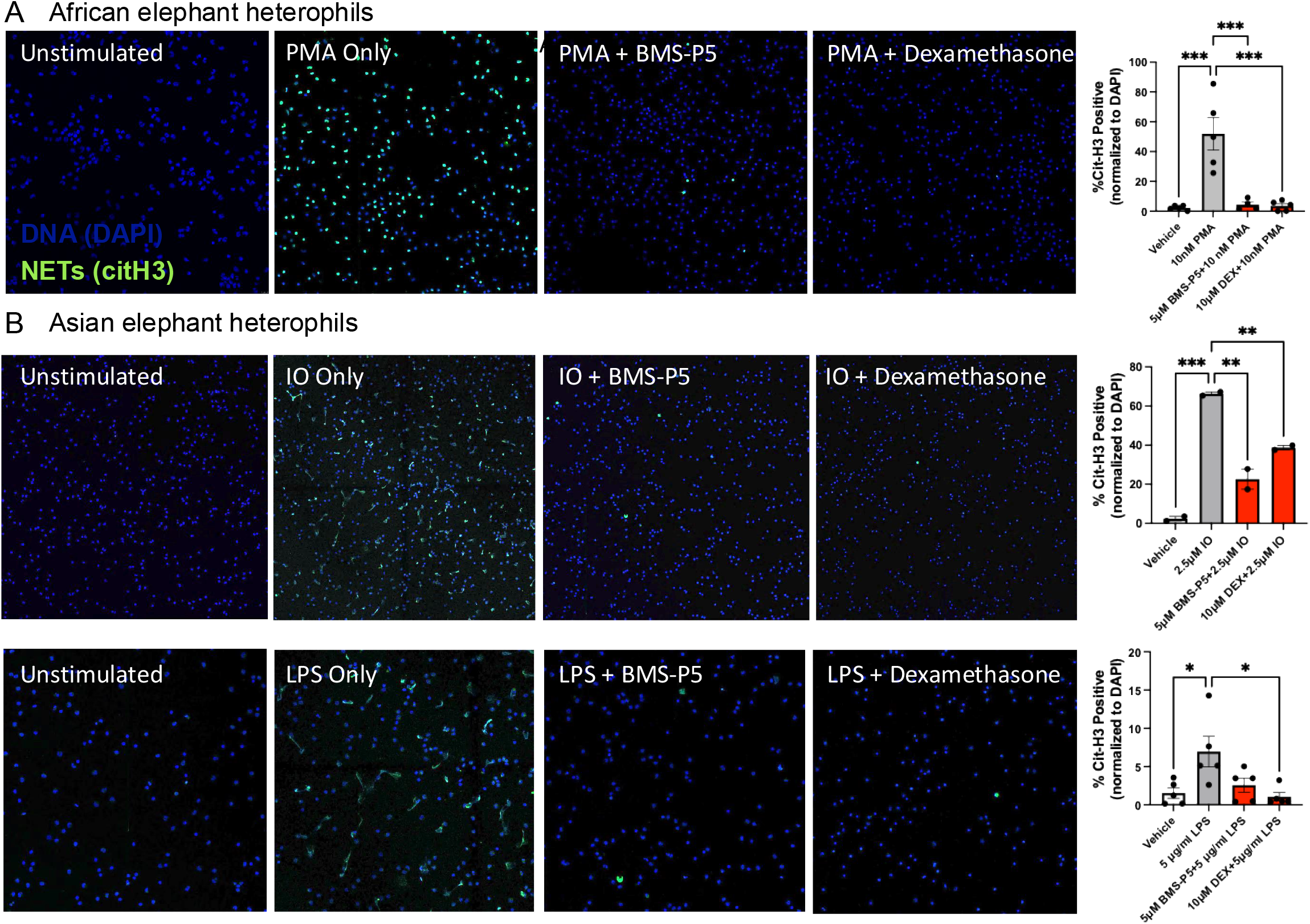
PMA, ionomycin, and LPS stimulated NET release from elephant heterophils were inhibited by BMS-P5 and dexamethasone. A. Representative images and quantification of African elephant heterophils treated with PMA after pretreatment with either vehicle, BMS-P5, or dexamethasone. Dots on graph represent two individual elephants each tested on two different days. B. Representative images and quantification of Asian elephant heterophils treated with ionomycin (IO) or LPS after pretreatment with either vehicle, BMS-P5, or dexamethasone. NETs were quantified and graphed. Dots on graphs represent different individual elephants tested. *p<0.05, **p<0.01, ***p<0.0005

**Figure 7:**
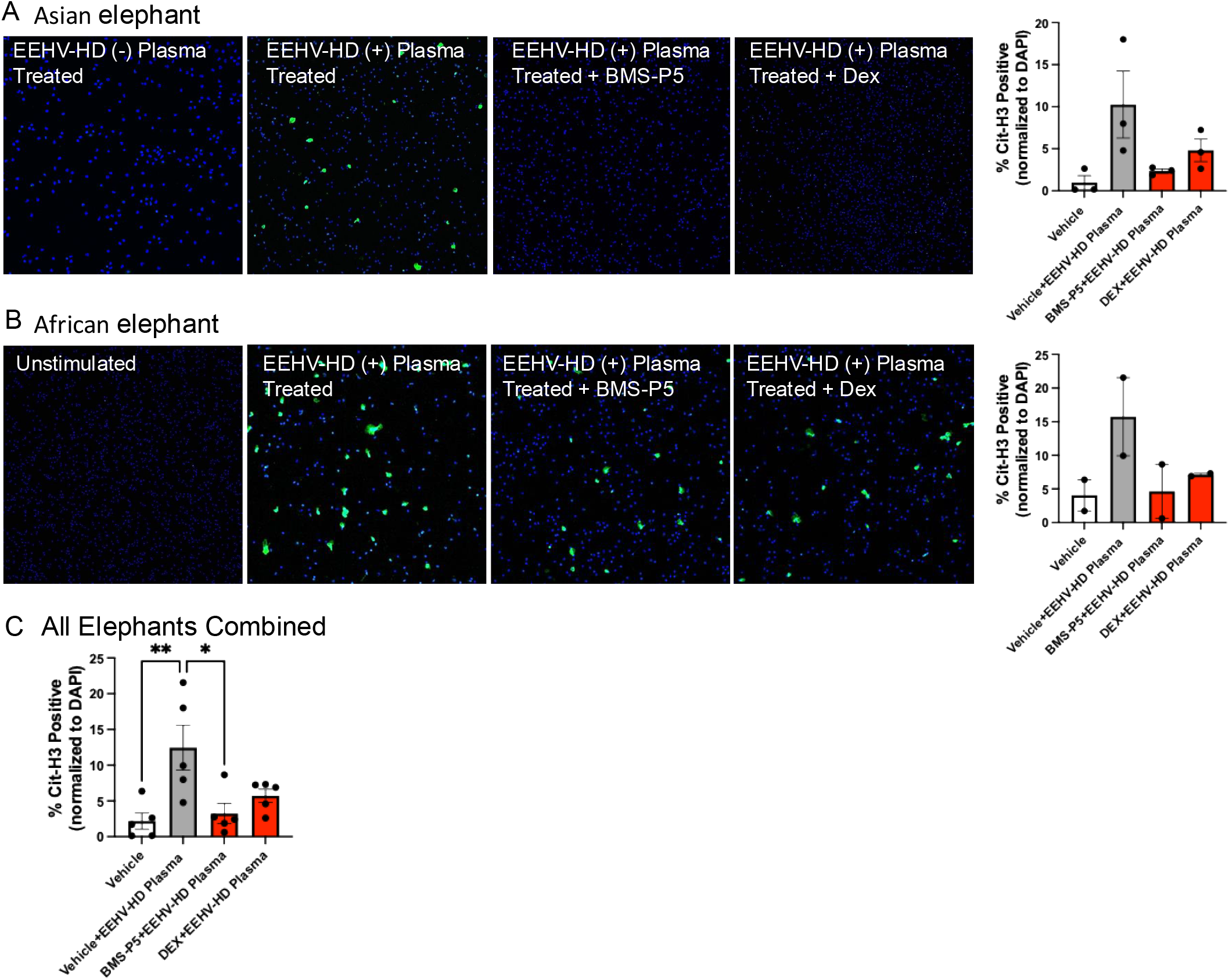
EEHV-HD positive plasma stimulated NET release from elephant heterophils was inhibited by BMS-P5 and dexamethasone. A. Representative images and quantification of Asian elephant heterophils treated with plasma from an EEHV-HD affected elephant after pretreatment with either vehicle, BMS-P5, or dexamethasone. The control in the first box is heterophils treated with EEHV-HD negative plasma. B. Representative images and quantification of African elephant heterophils treated with plasma from an EEHV-HD affected elephant after pretreatment with either vehicle, BMS-P5, or dexamethasone. The control in the first box is unstimulated heterophils. NETs were quantified and graphed. Dots on graphs represent different individual elephants tested. C. Combined quantification of NET release and inhibition for all elephants tested. *p<0.05, **p<0.01

## Discussion

We report for the first time that heterophils isolated from the blood of Asian and African elephants form NETs in response to stimulation. Significant NET formation was also observed in tissues from elephants that succumbed to EEHV-HD, along with evidence of NETs within lesions that are likely microthrombi. These findings align with previous reports of disseminated intravascular coagulation (DIC) and microthrombosis in some EEHV-HD-affected tissues ^31,32^. Given the established role of NETs in the pathogenesis of viral diseases in other species ^49^, including virus-induced clotting ^40^, our results suggest that NETs likely contribute to the development and lethality of hemorrhagic disease caused by EEHV viremia. Importantly, we report that NET formation can be inhibited in elephant immune cells, potentially offering a new therapeutic target for this lethal disease that threatens the survival of elephant species.

NETs play a damaging role in inflammation-related diseases, and several therapeutic companies are actively developing approaches to inhibit their formation ^62^. Although NET-specific inhibitors are not yet FDA-approved, other existing therapeutics can impact NET formation either by nonspecifically suppressing the immune response or by degrading NETs post-formation. For example, dexamethasone has shown survival benefits in humans with severe COVID-19 potentially through NET inhibition ^63^. Additionally, in various models of sepsis, NET inhibition or degradation has been associated with improved survival ^54–57^. Therefore, one of our study aims was to evaluate whether NET release from elephant heterophils could be inhibited and if this approach could provide future potential treatment options for elephants with EEHV-HD. We found that a compound designed to inhibit PAD4 activity (BMS-P5) and a general immune suppressor (dexamethasone) were both effective at inhibiting NET release from Asian and African elephant heterophils *in vitro*, including heterophils stimulated with plasma from EEHV-HD-affected elephants.

Based on these findings and the role of NETs in virus-induced thrombosis in humans ^40^, we propose a model whereby EEHVs replicate in endothelial cells lining small blood vessels leading to cell lysis. This damage activates platelets and clotting factors, triggering an inflammatory response and subsequent NET release from heterophils. The NETs then exacerbate endothelial damage and contribute to microthrombi by interacting with red blood cells, fibrin, and platelets. The formation of microthrombi and further endothelial damage likely exacerbates bleeding caused by viral lysis of endothelial cells (**Figure 8**). This robust immune response, which results in fatalities more frequently in Asian than African elephants with EEHV-HD, aligns with our earlier genomic studies that demonstrated selection of TNF-alpha response pathways in the evolution of Asian more than African elephants ^64^.

**Figure 8:**
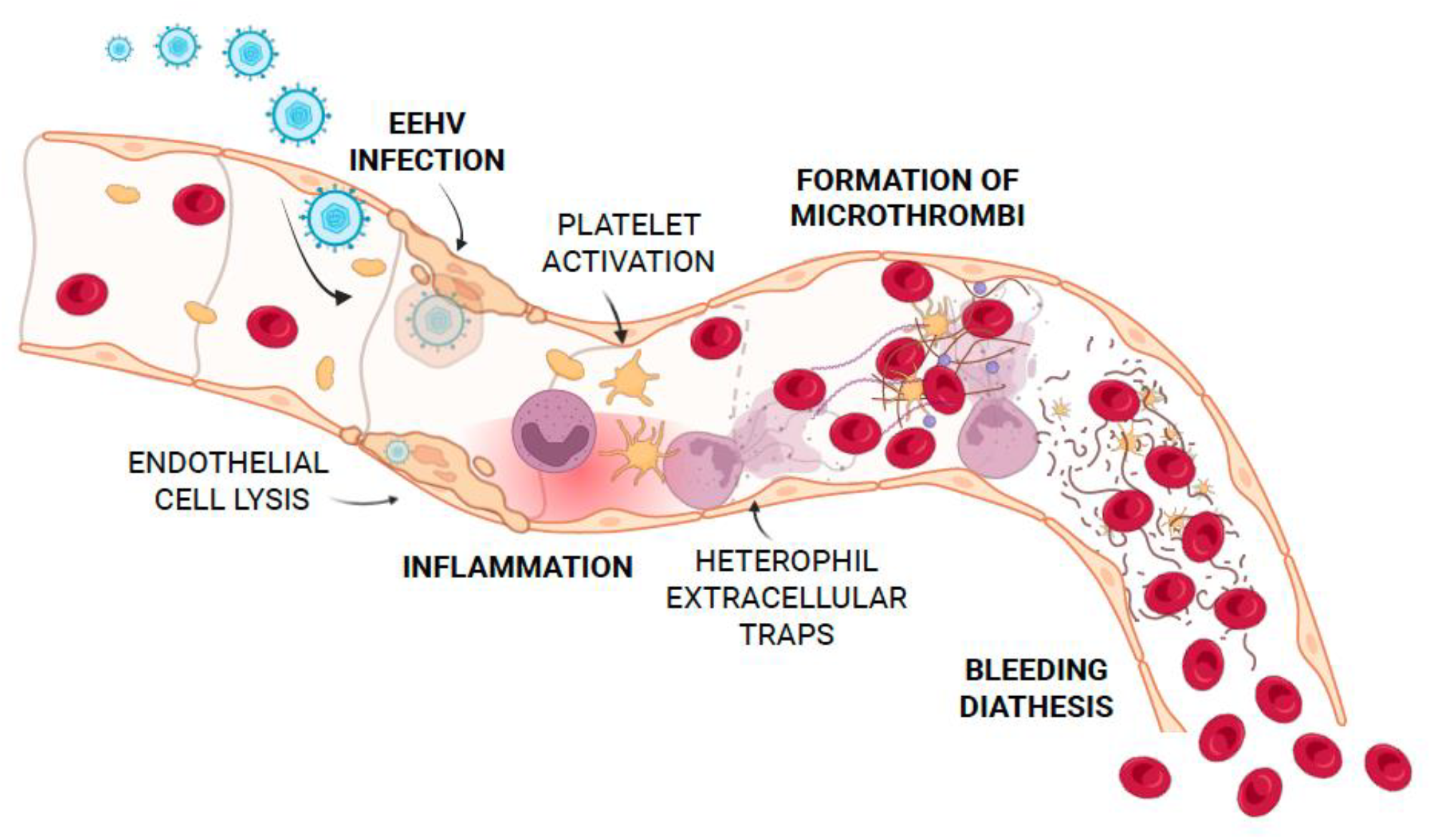
Model of EEHV-HD induced NET release and downstream immunothrombosis.

Our findings support NET inhibition as a new therapeutic target for EEHV-HD to decrease inflammation and impact clot formation, which potentially could improve outcomes of sick elephants. Inhibiting NET formation or degrading existing NETs has been shown to improve survival in animal models of DIC and sepsis ^54–57^. Steroid drugs like dexamethasone can inhibit NET formation by suppressing inflammatory transcription factors ^65^. In our study, dexamethasone effectively inhibited EEHV-HD positive plasma induced NET formation in isolated elephant heterophils, suggesting potential benefits for elephants with EEHV-HD. However, the broader immunosuppressive effects of dexamethasone should be carefully considered, as it may also suppress other beneficial immune responses. Case reports have described the use of dexamethasone alongside other supportive therapies in EEHV-HD-affected elephants, often as part of a comprehensive treatment plan including fluid therapy, antiviral medications, antibiotics, and supportive care ^22,66^. In humans, effective dosage and duration of dexamethasone treatment vary based on disease severity and response to treatment. This variability may also be the case in elephants. In some EEHV-HD cases, dexamethasone administration was associated with clinical improvement such as fever resolution, edema reduction, and stabilization of vital signs ^67^. However, treatment outcomes remain variable in elephants suffering from EEHV-HD and not all elephants will respond favorably to treatment, regardless of dexamethasone use.

While dexamethasone may alleviate inflammation, its use in EEHV-HD remains controversial among veterinarians due to concerns about potential adverse effects and its potential impact on viral replication. For instance, dexamethasone has been shown to increase the replication of cytomegalovirus, a herpesvirus that infects humans ^68^. Whether dexamethasone affects EEHV replication remains unknown, partly due to the challenges of culturing EEHV in vitro ^69^. The potential impact of dexamethasone on EEHV replication could be discovered by measuring viral titers before and after treatment in sick elephants. Ultimately, collecting and documenting clinical and laboratory data from elephants treated with and without dexamethasone will help clarify whether this drug offers a survival benefit in the context of EEHV-HD along with the risks of EEHV replication. In human viral infections, the timing and dosage of dexamethasone administration are critical to optimizing outcomes ^70^, suggesting that similar considerations will be necessary for elephants with EEHV-HD. Further research is required to determine the optimal dose, timing, and route of administration of dexamethasone to effectively decrease NET formation, vascular damage, and immunothrombosis without exacerbating other pathogenic aspects of the disease.

In addition to dexamethasone, we found that a commercially available NET inhibitor (BMS-P5) blocks EEHV-HD plasma induced NET formation. While this drug is not FDA approved and its future clinical development remains unclear, new therapeutic approaches to inhibit NET formation without affecting other immune functions are in active development for treating various inflammatory diseases, including DIC-induced thrombosis. Studies in mice demonstrated that blocking NET formation significantly improves survival in models of hemorrhagic disease, sepsis, stroke, and other conditions with similar symptoms of EEHV-HD ^54–56,71^. Once NET-specific inhibitors become clinically available, they should be carefully tested for efficacy in the context of EEHV-HD.

An FDA-approved approach that potentially mitigates the harmful effects of NETs is recombinant DNase I (dornase alpha or Pulmozyme), which digests NETs after their release. Pulmozyme is primarily used to treat cystic fibrosis and is administered via inhalation ^72,73^. During the COVID-19 pandemic, aerosolized DNase I was used with partial success to block fatalities from NET-associated acute respiratory distress syndrome ^40,74,75^. While lung administration may not benefit other organs affected by EEHV-HD, DNase has been explored in animal models using alternative delivery methods ^76^ and is in clinical trials for sepsis for intravenous administration (IdealSEPSIS1, NCT05453695). In addition, rectally administered DNase has shown a significant survival benefit in a mouse model of sepsis ^54^. This novel adaptation could be an approach worth investigating in EEHV-HD. Another potential therapeutic to consider testing in elephants is disulfiram, a drug that blocks acetaldehyde (ALDH) which may also disrupt NETs ^77^.

In conclusion, we offer the first demonstration of NET release in elephant heterophils and their potential contribution to EEHV-HD, a deadly coagulopathy in elephants. A thorough understanding of the interplay between viral infection and immune response is essential to improve outcomes for elephants affected by EEHV-HD. Ongoing efforts to characterize disease progression, facilitated by the North American EEHV Advisory Group ^21^, underscore the importance of collaboration among researchers, veterinarians, elephant care experts, and pathologists worldwide. Through continued research, cooperation, and careful analysis, these research collaborations can uncover effective treatment strategies to combat this devastating disease, ultimately helping to ensure the survival of elephant species.

## Acknowledgements

We would like to thank Deborah Olsen and Sara Conley at the International Elephant Foundation for their support of this project. We would like to thank David Gillespie and Randy Jensen for kindly allowing us to use their microscope (Olympus BX41). We thank Christian Con Yost and Mark Cody at the University of Utah for sharing their expertise related to measuring NET formation. We thank the University of Utah Cell Imaging Core for the use of their Zeiss AxioScan Z1 and for providing their expertise to support this project. We thank the Baylor College of Medicine Integrated Microscopy Core. The Integrated Microscopy Core is supported by the Center for Advanced Microscopy and Image Informatics (CAMII) with funding from NIH (DK56338, CA125123, ES030285), and CPRIT (RP150578, RP170719). We would like to thank Aaron Bretones for administrative support and Matthew Buccilli for technical support. We thank Utah’s Hogle Zoo, Houston Zoo, Rosamond Gifford Zoo, Columbus Zoo, San Antonio Zoo, ABQ BioPark Zoo, Denver Zoo, Santa Barbara Zoo, Oklahoma City Zoo, Dickerson Park Zoo, and others for providing samples and for their tremendous dedication and tireless efforts to save elephants from succumbing to EEHV-HD.

This work was supported by Arizona Cancer Evolution Center Pilot Funding from NIH grant U54 CA217376 and University of Utah Department of Pediatrics Research Enterprise.

